# From Planning Stage To FAIR Data: A Practical Metadatasheet For Biomedical Scientists

**DOI:** 10.1101/2024.01.27.577552

**Authors:** Lea Seep, Stephan Grein, Iva Splichalova, Danli Ran, Mickel Mikhael, Staffan Hildebrand, Mario Lauterbach, Karsten Hiller, Dalila Juliana Silva Ribeiro, Katharina Sieckmann, Ronja Kardinal, Hao Huang, Jiangyan Yu, Sebastian Kallabis, Janina Behrens, Andreas Till, Viktoriya Peeva, Akim Strohmeyer, Johanna Bruder, Tobias Blum, Ana Soriano-Arroquia, Dominik Tischer, Katharina Kuellmer, Yuanfang Li, Marc Beyer, Anne-Kathrin Gellner, Tobias Fromme, Henning Wackerhage, Martin Klingenspor, Wiebke K. Fenske, Ludger Scheja, Felix Meissner, Andreas Schlitzer, Elvira Mass, Dagmar Wachten, Eicke Latz, Alexander Pfeifer, Jan Hasenauer

**Affiliations:** Computational Biology, Life & Medical Sciences (LIMES) Institute, University of Bonn, Bonn, Germany; Developmental Biology of the Immune System, Life & Medical Sciences (LIMES) Institute, University of Bonn, Bonn, Germany; Institute of Pharmacology and Toxicology, University Hospital, University of Bonn, Bonn, Germany; Department of Bioinformatics and Biochemistry, Technical University Braunschweig, Braunschweig, Germany; Institute of Innate Immunity, University Hospital Bonn, University of Bonn, Bonn, Germany; Quantitative Systems Biology, Life & Medical Sciences (LIMES) Institute, University of Bonn, Bonn, Germany; Systems Immunology and Proteomics, Institute of Innate Immunity, Medical Faculty, University of Bonn, Bonn, Germany; Department of Biochemistry and Molecular Cell Biology, University Medical Center Hamburg-Eppendorf, Hamburg, Germany; Department of Internal Medicine I, Division of Endocrinology, Diabetes and Metabolism, University Medical Center Bonn, Bonn, Germany; Chair of Molecular Nutritional Medicine, TUM School of Life Sciences, Technical University of Munich, Freising, Germany; Immunology and Environment, Life & Medical Sciences (LIMES) Institute, University of Bonn, Bonn, Germany; EKFZ—Else Kröner-Fresenius Center for Nutritional Medicine, Technical University of Munich, Freising, Germany; Immunogenomics & Neurodegeneration, German Center for Neurodegenerative Diseases (DZNE), Bonn, Germany; PRECISE, Platform for Single Cell Genomics and Epigenomics at the German Center for Neurodegenerative Diseases and the University of Bonn, Bonn, Germany; Department of Psychiatry and Psychotherapy, University Hospital Bonn, Bonn, Germany; Institute of Physiology II, Medical Faculty, University of Bonn, Bonn, Germany; School for Medicine and Health, Faculty of Sport and Health Sciences, Technical University of Munich, Munich, Germany; ZIEL Institute for Food & Health, Technical University of Munich, Freising, Germany; Department of Internal Medicine I - Endocrinology, Diabetology and Metabolism, Gastroenterology and Hepatology, University Hospital Bergmannsheil, Bochum, Germany; Experimental Systems Immunology, Max Planck Institute of Biochemistry, Martinsried, Germany; PharmaCenter Bonn, University of Bonn, Bonn, Germany; Helmholtz Center Munich, German Research Center for Environmental Health, Computational Health Center, Munich, Germany

## Abstract

Datasets consist of measurement data and metadata. Metadata provides context, essential for understanding and (re-)using data. Various metadata standards exist for different methods, systems and contexts. However, relevant information resides at differing stages across the data-lifecycle. Often, this information is defined and standardized only at publication stage, which can lead to data loss and workload increase.

In this study, we developed Metadatasheet, a metadata standard based on interviews with members of two biomedical consortia and systematic screening of data repositories. It aligns with the data-lifecycle allowing synchronous metadata recording within Microsoft Excel, a widespread data recording software. Additionally, we provide an implementation, the Metadata Workbook, that offers user-friendly features like automation, dynamic adaption, metadata integrity checks, and export options for various metadata standards.

By design and due to its extensive documentation, the proposed metadata standard simplifies recording and structuring of metadata for biomedical scientists, promoting practicality and convenience in data management. This framework can accelerate scientific progress by enhancing collaboration and knowledge transfer throughout the intermediate steps of data creation.

## 1 Introduction

Collaboration along with the open exchange of techniques, protocols and data are the backbone of modern biomedical research^1^. Data usage and retrieval requires the structured collection of information, such as study design, experimental conditions, sample preparation and sample processing, on the performed measurements. This information is generally referred to as metadata which grows along the research data-lifecycle (Fig. 1A), from planning to its final storage alongside publication.^2–6^. There is a growing consensus among researchers, journals and funding agencies that data should adhere to the principles of being findable, accessible, inter-operable and reusable (FAIR). The adherence to these FAIR data principles^7^ requires metadata^8,9^ and benefits from the use of trustworthy digital repositories. Trustworthiness is marked by Transparency, Responsibility, User focus, Sustainability and Technology (TRUST)^10^.

**Figure 1.**
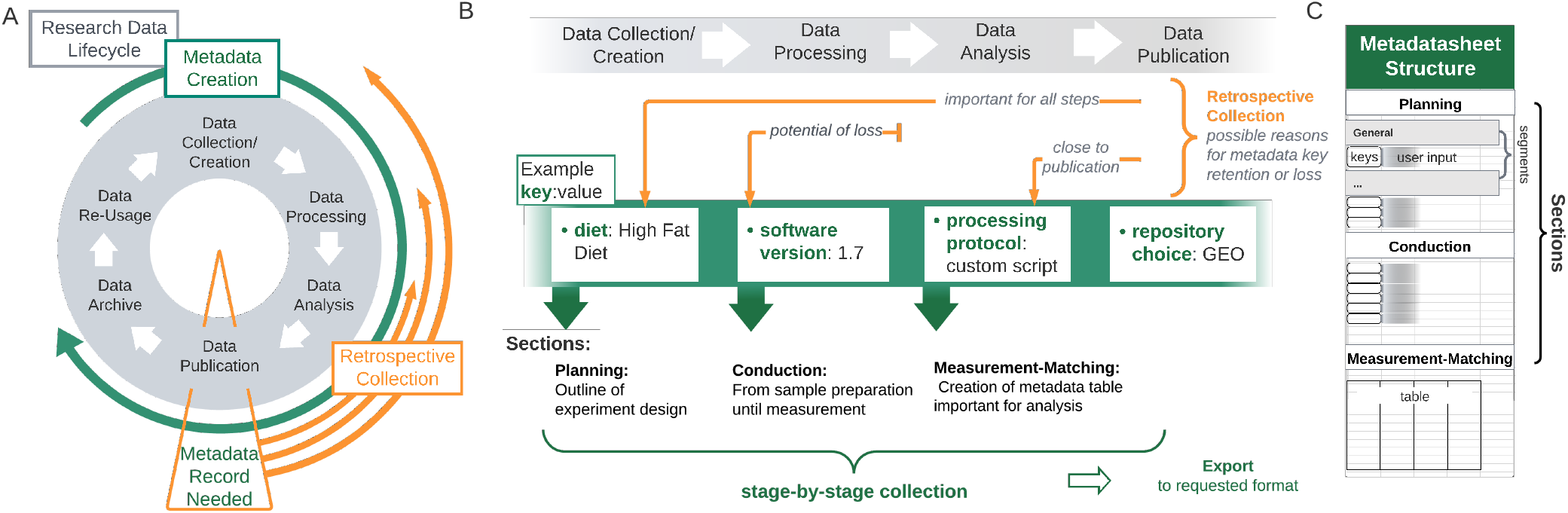
Alignment of Metadata Lifecycle with the Research Data-Lifecycle. **(A)** Metadata is created alongside the research data creation, however, often only gathered at the point of publication when it is requested from, e.g., repositories. **(B)** Through the retrospective collection, recovering necessary metadata items can be challenging. When not generally recorded, only metadata-items of immediate interest for the next step or for data analysis will be available, other items that ensure FAIRness, e.g., the software version of a specific program, might not get recorded at all and will hence be lost.**(C)** The structure of the proposed Metadatasheet is defined by its sections, which further encompass segments. Within each segment user input is required, which can be of different forms, e.g., values to keys or table entries.

Repositories are subdivided into cross-discipline and domain-specific categories. Cross-discipline repositories intentionally do not impose any requirements on format or size to allow sharing without boundaries. Domain-specific repositories in the field of biomedicine such as BioSample and GEO^11^, maintained by the National Center for Biotechnology Information (NCBI), or PRIDE^12^ and BioModels^13,14^, maintained by European Bioinformatics Institute (EBI), impose requirements during submission in form of data and metadata standards. Standards often make use of controlled vocabularies and ontologies to ensure consistency and comparability. Controlled vocabularies, consisting of standardized terms, describe requested characteristics and keys^5^, while ontologies, such as the Gene Ontology (GO)^15^, establish structured frameworks for depicting relationships between entities, fostering comprehensive and searchable knowledge structures. Current metadata standards can be divided into two categories. First, comprehensive high-level documents that are often tailored to specific use cases. These documents primarily consist of lists of requested terms or guidelines, often interconnected with corresponding ontologies. For instance, ARRIVE (Animal Research: Reporting of In Vivo Experiments) provides a checklist of information to include in publications of *in vivo* experiments^16^ or MIRIAM (minimum information requested in the annotation of biochemical models)^17^ standardizes the curation of biochemical models including their annotations. Second, there are structured metadata standards supplied and requested by respective repositories. Irrespective of the suitable metadata standard, it is common to adhere to requested standards at the stage of data publication evoking a retrospective collection (Fig. 1A). Necessary information resides at all stages of the data-lifecycle and may involve different responsible individuals, thereby rendering the retrospective metadata collection resource-intensive.

Despite the existence of various metadata standards in biomedical sciences and widespread recognition of the relevance of metadata, a practical issue persists: the absence of a dedicated metadata standard that effectively and with low burden directs researchers in capturing metadata along the data-lifecycle without loss of information, ensuring FAIRness during and after the experiment (Fig. 1B). Thus, we propose a metadata standard tailored for wet-lab scientists mirroring the phases of the biomedical research lifecycle, offering transferability across distinct stages and among diverse stakeholders.

The proposed standard, further referred to as Metadatasheet, is embedded in a macro-enabled Excel workbook, further referred to as Metadata Workbook. The Metadata Workbook offers various usability features, such as automation, integrity checks, extensive documentation, usage of templates, and a set of export functionalities to other metadata standards. By design, the proposed Metadatasheet, accompanied by the Metadata Workbook, naturally allows stage-by-stage collection, embodying a paradigm shift in metadata collection strategies, and promoting the efficient use of knowledge in the pre-publication phase.

## 2 Results

### 2.1 The Metadatasheet is based on comprehensive interview of biomedical researchers

Metadata information consists of a set of characteristics, attributes, herein named keys, that intend to provide a common understanding of the data. Example keys are experimental system, tissue type, or measurement type. Accordingly, the Metadatasheet is built upon requested keys gathered from comprehensive interviews of research groups and systematical collection from public repositories. In the initial phase more than 30 experimental researchers from the biomedical sciences participated, who were from two consortia focusing on metaflammation (https://www.sfb1454-metaflammation.de/) and metabolism of brown adipose tissue (https://www.trr333.uni-bonn.de/). The participating researchers reported common general keys as well as diverse experimental designs covering 5 major experimental systems and 15 common measurement techniques, each accompanied with their specific set of keys. To refine and enhance the set of metadata keys, we engaged in iterative consultations with biomedical researchers. In parallel, we systematically collected relevant keys from four popular public repositories, namely NCBI^18^, GEO^11^, the Metabolomics Workbench^19^ and the PRIDE^12^ database. Moreover, expected input, summarized under the term ‘controlled vocabulary’, for all keys needed to be specified. From second iteration on, specifications of the controlled vocabulary, as well as the set of keys, were improved based on researchers’ feedback. The comprehensive key and controlled vocabulary collection process revealed the dynamic, unique and growing requirements of different projects, in terms of values within the controlled vocabulary and performed measurements. Those requirements lead to the choice of allowing customisation and expansion of key sets and controlled vocabulary as an integral part of the Metadatasheet. To handle the dynamic and adaptable nature of the Metadatasheet, it was embedded within a reactive framework with additional functionalities, the Metadata Workbook.

In the following, the overall concept and design of the Metadatasheet is introduced, afterwards key aspects of the Metadata Workbook are highlighted. The results section concludes with an example Metadatasheet generated by the Metadata Workbook.

### 2.2 The Metadatasheet design follows and allows metadata recording along the data-lifecycle

The proposed Metadatasheet is organized into three main sections: ‘planning’, ‘conduction’ and ‘measurement-matching’ section. These sections mirror the stages of the data-lifecycle and align with the general experimental timeline (Fig. 1B). The analogous top-to-bottom structure allows sequential metadata recording acknowledging the continuous growth of metadata. Each section further subdivides into segments, which hold the keys, that need to be specified by the user through values. The segmentation aims to group keys into logical units, that are likely provided by a single individual. This grouping enables the assignment of responsible persons, resulting in a clear emergent order for data entry if multiple persons are involved. Moreover, within a section the segments are independent from each other allowing also parallel data entry.

Metadatasheet keys can be categorized based on the form of the expected input. First, providing a single value (key:value pair), e.g. the analyzed ‘tissue’ (key) originates from the ‘liver’ (value). Second, filling tables, whereby the row names can be interpreted as keys but multiple values need to be provided (one per column). Third, changing a key:value entry to a table entry by the keyword ‘CHANGES’. If the keyword is supplied as a value, the respective target key changes from key:value pair to a table entry. The switch of form allows data entries to be minimal if sufficient or exhaustively detailed if needed. This flexible data entry minimizes the need of repetition gaining easier readability but allows recording fine-grained information whenever needed.

Required values can be entered in form of controlled vocabulary items, date-format, free text including numbers or filenames. Filenames are a special type of free text and specify additional resources where corresponding files are either expected within the same directory as the Metadatasheet itself or given as relative path. Suitable form of values is naturally determined by the key, e.g., ‘Date’ is of date format, ‘weight’ is of number format and ‘tissue’ of discrete nature to be selected from the controlled vocabulary. The format choice is constraining the allowed values. Providing such input constraints to each key, allows harmonization of metadata. Harmonization enables machine readability which is a starting point for further applications.

A single Metadatasheet captures the combination of an experimental design and a measurement type, as those come with a distinct set of keys, also referred to as dependent keys. An experimental design is here defined as a specific experimental system exposed to a contrasting setting. Within the Metadatasheet five contrasting settings, herein named comparison groups, are set: ‘diet’, ‘treatment’, ‘genotype’, ‘age’, ‘temperature’ and ‘other’ (non-specific). Experimental designs exhibit a range of complexities, they can span multiple comparison groups such as different treatments exposed to different genotypes, while each group can have multiple instances such as LPS-treatment and control-treatment.

The varying complexity in experimental designs is reflected in the Metadatasheet structure. This reflection is achieved through hierarchies, organized into up to three levels. The top-level keys are mandatory, while the inclusion of other-level keys depends on design’s complexity. Present hierarchies within the samples are also important to consider for statistical analysis. Hierarchies emerge, if the sample is divided into subsamples prior to the measurement. For instance, if the experimental system involves a mouse with two extracted organs for measurement, the relation to the sample should be specified. Moreover, subsamples are also present when measurements where conducted on technical replicates of the extracted sample. The Metadatasheet accommodates up to two levels of sample partitioning. By leveraging a hierarchical structure, details are displayed only when necessary, avoiding unnecessary intricacies. Moreover, relationships of the measured samples can be recorded, enhancing clarity.

To ensure coherence between a samples’ actual measurement data and recorded metadata, it is crucial to link them accurately by an unique personal ID. To guide through matching and prevent mismatches, we have designed the Measurement-Matching section to summarize essential information and focusing on differences between samples. This information includes their association with an instance of a comparison group, the number of replicate, and the presence or absence of subsamples. If subsamples are present, they are organized in a separate table, referencing their higher, preceding sample. Careful recording also involves specified covariates. They are expected at the lowest level, the measurement level, and must be carefully matched to the correct ID within the set of replicates within a comparison group instance.

The inherent innovative force within the research community risk to hit boundaries of anything predefined, here, particularly evident in controlled vocabulary and dependent keys. Those predefined sets come as additional tables, associated with the Metadatasheet. Subsequently, the resources of the Metadatasheet require an ongoing commitment to be extended and further developed. The separation of the Metadatasheet and its resources also allows the creation of group-specific subsets of controlled vocabulary. This feature proves helpful when a group wants a more constrained set of controlled vocabulary, e.g., using ontologies and respective value specifications. The group-specific validation should be a subset of the overall validation.

The Metadatasheet design aligns with the data-lifecycle to allow analogous metadata recording. Presented design choices allow to adopt to various settings biomedical researchers are confronted with and thereby provide a high degree of flexibility.

### 2.3 The implementation of the Metadatasheet, the Metadata Workbook, enhances user experience by automation, integrity checks, customisation and export to other formats

Gathering the diverse resources, specifically the Metadatasheet, the validation and dependent fields resources, we created an Excel Workbook including all of those sheets. To promote usage through user-friendliness, dynamic adaption and automation, we further introduced Excel macros (a set of custom functions) resolving to a macro enabled Excel workbook, called the Metadata Workbook. This Metadata Workbook is designed to guide the Metadatasheet application while providing automation whenever possible. Advancements through the implementation include specifically the ability of automatic insertion of the depending keys, enhancements to user experience, easy expansion and updating of the controlled vocabulary, the option to use templates, automatic checks of data input validity as well as the export of the Metadata Workbook to other formats allowing long-term storage. Crucial advancements are explained in more detail in the following.

The Metadata Workbook creates tailored Metadatasheets for common biomedical experimental systems and measurement techniques. Those segments come with their unique set of dependent keys and therefore change between individual Metadatasheets. Static sheets result therefore in a high amount of sheets. The Metadata Workbook provides a dynamic solution reducing different requirements to a single Metadata Workbook that needs to be handled. The dependent, inserted keys, can be extended, but not changed, by adding values to the respective column within the dependent field sheet. The new addition is automatically added to the validation sheet, holding the controlled vocabulary. For new additions, the key’s input constraints can be changed. These features enable flexibility through expansion, allowing to match current and future research contexts.

The Metadata Workbook employs various features to enhance user experience and convenience while facilitating to capture simple to advanced setups of an experiment: sections of the sheet collapse, such as second levels of hierarchical segments, if not applicable; DropDown menus based on the provided controlled vocabulary enrich value fields, facilitating ease of selection. Furthermore, visual cues notify users in several situations: any segment where the structure deviates from the typical key:value format to adopt to a tabular arrangement is highlighted automatically; text-highlighting is used to mark mistakes, e.g., if input values for key fields do not align with the controlled vocabulary. Altogether, Metadata Workbook provides a user-friendly environment to guide users to record metadata.

Disruptive redundancy across and within the proposed Metadatasheet is tackled within the Metadata Workbook. Redundancy across Metadatasheets occurs if multiple studies are conducted in the same context, with similar designs, systems or experimental techniques. To reduce redundancy and prevent mistakes from copying and pasting, existing Metadatasheets can serve as templates. All information from the first two sections (planning and conduction) are exported from an uploaded Metadatasheet. Upon upload, users only need to update the ID information in the Measurement-Matching section for the new setting. This exception prevents not updating these crucial IDs. Redundancy within a single Metadatasheet occurs while providing the ‘final groups’ as well as the table within the Measurement-Matching section at the beginning of section two and three, respectively. The Metadata Workbook provides ‘generate’ buttons to produce both those tables automatically. Hence the first ‘generate’ button creates all possible combinations based on the Planning section, while the measurement-matching table is generated based on the Conduction section. To maintain structural integrity, the Metadata Workbook requires a sequential input of the sections, the generate buttons prevent from violations by evoking an error if input in the preceding section is invalid. The ‘generate’ functionalities remove through automation, again, the need for copy paste actions and redundant actions for the user.

Upon the completion of the Metadata Workbook, it can be exported to various formats serving different objectives, such as compatibility with open-source software, long-term storage through TRUST repositories and minimization of work by don’t repeat yourself (DRY) principles^20^. Compatibility of the Metadata Workbook with open-source software, like LibreOffice, is facilitated by the export option to a simple Excel (xlsx filetype) file while simultaneously removing any associated functionalities. Notably, a unique identifier is automatically assigned upon export. Providing metadata represents a critical prerequisite before uploading data to repositories or publication. Repositories normally adhere to their distinct metadata standards. Some offer submission tools featuring user interfaces, e.g. MetabolomicsWorkbench. Conversely, others like GEO or NCBI require the manual completion of an Excel table. For both repositories, export capabilities have been added to transform the Metadata Workbook compliant with the repositories’ requirements. The proposed structure covers all mandatory fields from the major repositories. These export functionalities reduce the hours spend on reformatting to meet different requirements and are a crucial step towards DRY principles within the metadata annotation procedure. Further, a converter is provided that turns the proposed structure, given as an exported xlsx file, to an object, commonly used as input to data analysis. The converter, applicable to omics-data and associated metadata, returns an R object called SummarizedExperiment^21^. The SummarizedExperiment object can be easily shared and lays the foundation for a plethora of standardized bioinformatic analyses within R. The object contains all available metadata from previous data-lifecycle stages limiting issues due to missing information, like unmentioned covariates.

In essence, the introduced implementation results in a macro-enhanced Excel Workbook, the Metadata Workbook, with advanced functionalities that chooses the appropriate keys, enhances user experience with color cues and automation while maintaining data integrity.

### 2.4 Showcase and application of the Metadatasheet demonstrate its use in recording metadata and subsequent data analysis

To assess the suitability and adaptability of the designed Metadatasheet we asked researchers from 40 different groups to gather and transfer their metadata in this format. The initiation of capturing standardized metadata alongside the data generation process has made a range of practical applications possible, yielding multiple advantages within the consortia. The versatility of the proposed structure is demonstrated by a curated collection of sheets (Table 1), each accompanied by a concise description of the study’s setting. The provided selection encompasses various measurement types and differing experimental systems. The experimental designs within this selection range from straightforward setups to nested designs as well as two-way comparisons. For all complete Metadatasheets, see Supplementary Material. As the Metadatasheet records metadata from the start of the data-lifecycle, some measurement data in certain showcases is not included here due to its non-disclosure status before publication.

**Table 1.** Overview of curated collection of completed Metadatasheets, which can be found in the Supplementary Material.

| Measurement Type | Experimental System | Experimental Design | Notes | Provider |
| --- | --- | --- | --- | --- |
| bulk RNASeq | mouse | 6 diets | part of collection; manuscript example | I. S., H. H., E. M. |
| metabolomics (13C glucose) | human-derived | 2 treatments x 2 timepoints | time dependent timeline | M. L., K. H. |
| bulk proteomics | mouse | 2 others | stress-treatment; with drop out | A. K. G. |
| bulk proteomics | human-derived | 4 others | athlete groups; nested design (subsamples time) | A. S. A., F. M., S. K., H. W. |
| 16S rRNA Seq | rat | 4 others | bariatric surgery or fecal microbiota transfer | V. P., A. T., W. K. F. |
| indirect calorimetry | mouse | 2 genotypes | nested design (subsamples time) | S. H., A. P. |
| FACS | patient | 3 others | disease stages | J. Y., A. Sch. |
| single-cell RNASeq | mouse | 4 diets x 2 genotypes | time dependent timeline | Y. L., M. B. |
| single-nucleus RNASeq | mouse | 2 genotypes | nested design (subsamples tissue) | K. K., T. F. |
| bulk lipidomics | mouse | 2 diets x 2 genotypes | 2-fold comparison and nested design (subsamples tissue) | J. Be., L. Sch. |
| lipolysis measurement | cell-line | 9 treatments | nested design (subsamples technical replicates); well plate measurements | D. Ra., A. P. |
| UPLC-UV | cell-line | 6 treatments | nested design (subsamples technical replicates) | M. M., A. P. |
| FRET | cell-line | 1 treatment (timeseries) | timeseries involves the consecutive treatment with drugs | D. Ra., A. P. |
| Histology | mouse | 2 genotypes | multiple covariates given | R. K., K. S. D. W. |

In the following, a single Metadatasheet from the showcase collection is highlighted, which has been created with the Metadata Workbook. The picked Metadatasheet for demonstration encompasses one of the datasets associated with the study of developmental programming of Kupffer cells by maternal obesity^22^. The associated data is deposited on GEO and are accessible through GEO Series accession number GSE237408 (https://www.ncbi.nlm.nih.gov/geo/query/acc.cgi?acc=GSE237408).

### Example Planning Section

The Metadatasheet starts with the Planning section which captures all information already available during the conceptualization of an experiment. The section is subdivided into the segments ‘General’, ‘Experimental System’ and ‘Comparison groups’ (Fig. 2). The requested information in ‘General’ (Fig. 2A) includes personal information, the title of the project as well as the specification whether the sheet is part of a collection of multiple related Metadatasheets. Collections allow users to link individual Metadatasheets from the same project to spread awareness of such connections, in this example linking multiple datasets associated with the same project. ‘Experimental System’ segment provides automatically predefined keys (dependent fields sheet) after the selection within the Metadata Workbook, for example, ‘line’ and ‘genotype’ information will be needed upon selecting ‘mouse’ (Fig. 2B).

**Figure 2.**
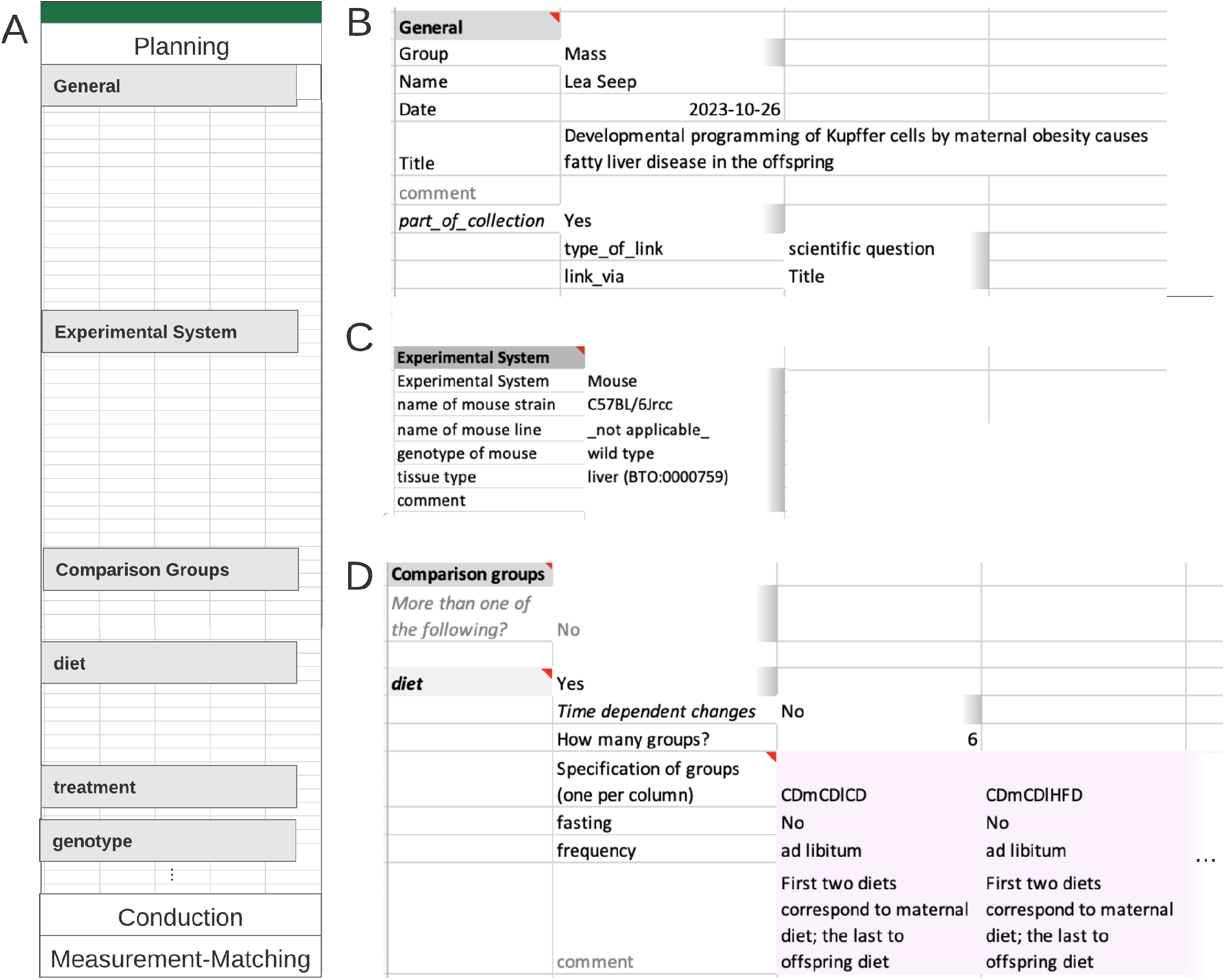
Example of an instance of the Planning section. **(A)** Overview Planning section. **(B)** General segment contains contact information and general project information in form of key:value pairs; on its second level, linked Metadatasheets can be specified. **(C)** The experimental system segment is requesting keys dependent on the value given to key ‘Experimental System’. **(D)** Comparison group segment; here further the only comparison group is ‘diet’. defined through diet (other comparison group options as treatment etc. not shown). As six groups are requested by the user a table is present with six columns (only two shown). Information per specified group is expected column-wise. Note that the full Metadatasheet of this example can be found in Supplementary Material.

The ‘Comparison groups’ segment (Fig. 2C) specifies the experimental design linked to the current research question. The experiment design for each comparison group involves two levels: broader comparison group, here ‘diet’ and details for each instance within the broader comparison group. Users are not restricted to a single comparison group. At the second level, details for each chosen comparison group are entered. Here, 6 different groups with varying diet schemes were studied. The established feeding scheme is unique within the consortia, those special requirements were easily added to the controlled vocabulary for ‘diet’ with the Metadata Workbook, leveraging on its adaptability.

### Example Conduction Section

The Conduction section is divided into six segments and captures all information created during the experimental/ wet-lab phase. The section starts with the specification of the ‘final groups’ resulting form previously specified comparison groups. As diet is the only comparison group with six instances, the final groups resolve to those types (Fig. 3A). If multiple groups are planned, for example, if six diet groups and two genotype groups, 12 final groups would be present due to all combination possibilities. Within the Metadata Workbook those final groups are generated automatically, the user then defines the respective replicates.

**Figure 3.**
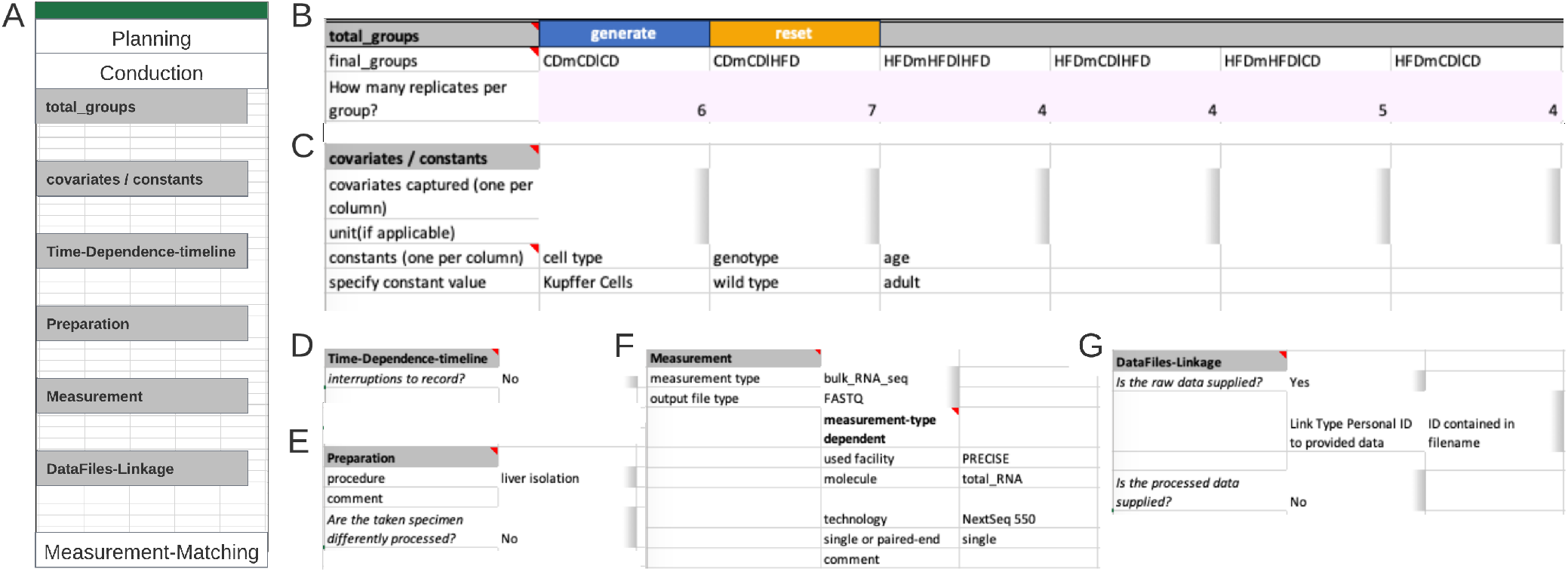
Example of an instance of the Conduction section. **(A)** Overview Conduction section. **(B)** The ‘total_groups’ segment expects all possible combinations of the comparison groups defined in the Planning section. Number of replicates belongs underneath each group. In the Metadatasheet implementation ‘final_groups’ are generated; pink color marks an expected table. **(C)** The segment covariates / constants requests respective specification including units. For constants the value is expected in place, whereas covariates values are expected within the measurement-matching table. **(D)** Time-Dependence-timeline segment collapses completely if not required. **(E)** Preparation segment expects the procedure that is required before the actual measurement. Here, the reference to either a fixed protocol, chosen from the controlled vocabulary or a filename is expected. The specified file is expected to be on the same level as the Metadatasheet in the filesystem. **(F)** The Measurement segment is requesting keys depending on the value given to key measurement type. **(G)** The DataFiles-Linkage segment specifies how to identify the correct measurement file given the subsequent (within the measurement matching section) specified personal ID. If there is no clear pattern one can choose keyword ‘CHANGES’ to promote filename specification to the measurement matching section. Note that the full Metadatasheet of this example can be found in Supplementary Material.

The segment ‘Covariates/Constants’, expects each constant or covariate to fill a single column with the respective suitable unit (table form). For clarification, a covariate refers to any additional variable or factor, beyond the main variables of interest (comparison groups), that is considered or observed in the experimental design. This could include factors such as age, gender, environmental conditions but also unusual colour of serum or day of preparation. Here, no covariate but the constants ‘cell type’ and ‘genotype’ were recorded, respective values, ‘Kupffer Cells’ and ‘wild type’ occupying a single columns each (Fig. 3B).

The next segment ‘Time-Dependence-Timeline’ is organized hierarchically. On the first level, one decides whether this segment is applicable, by answering if interruptions are present. The presence of an interrupted timeline is given, when the designated comparison group is to be augmented with temporal details that occurred during the experimental period. The second level distinguishes between two types of an interrupted timeline: ‘continued’ and ‘discontinued’. A ‘continued’ timeline is identified when temporal details are annotated. On the other hand, if the temporal details describe a change, such as a modification in treatment, it falls under the ‘discontinued’ type. For example, an interrupted timeline is present when a mouse undergoes several glucose tolerance tests during a contrasting diet setting (interrupted timeline type continued), or when a treatment consists of administering agent A for 24 hours followed by agent B for the next hours (interrupted timeline type discontinued). While not present at the example at hand, both types of interrupted timelines would require further details (Fig. 4A).

**Figure 4.**
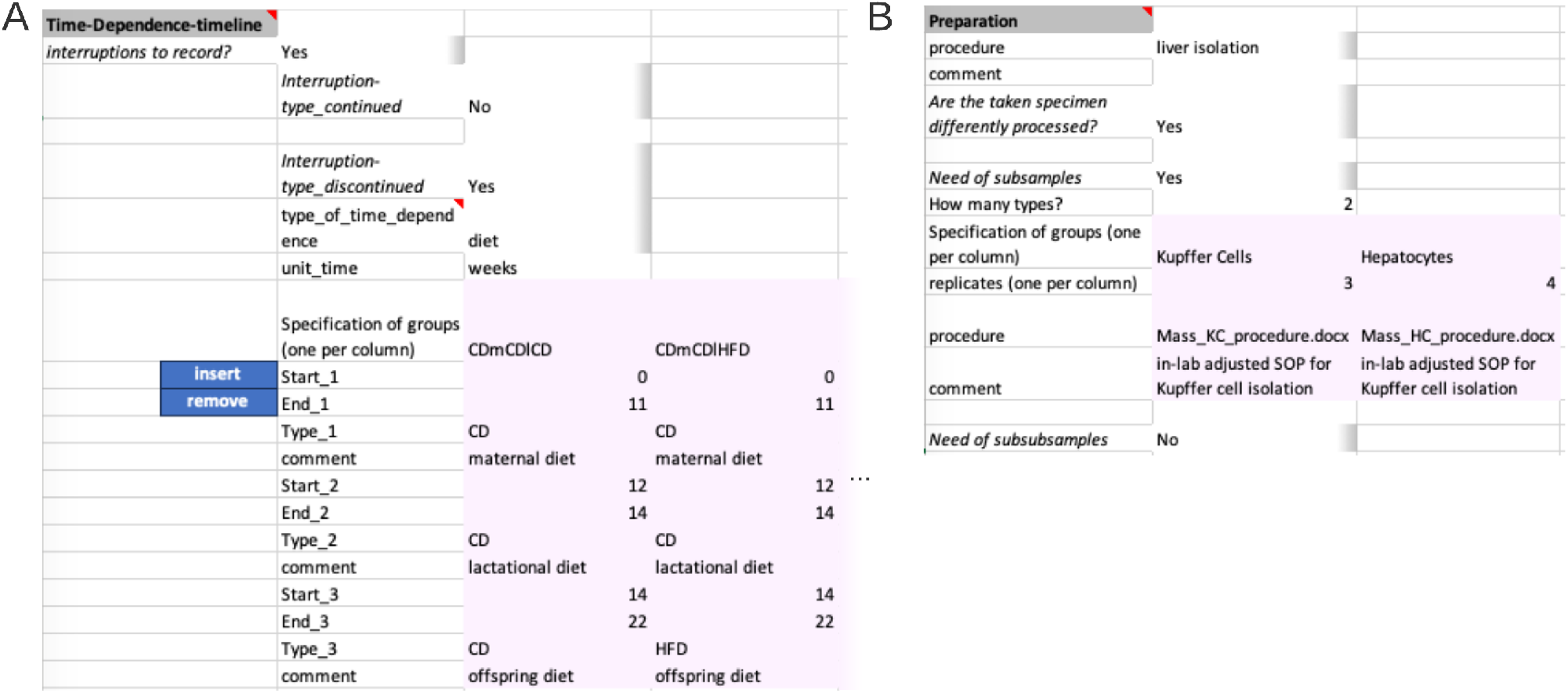
Advanced example of segments within the Conduction section. **(A)** Within the Time-Dependence Timeline segment, given comparison groups can be enriched with time dependent information on the second hierarchy level. One specifies which of the comparison groups is to be enriched with timeline information and the unit of time. Then, time-steps can be specified. Pink color marks the table, which needs to be filled. **(B)** Within the Preparation segment one can supply up to two divisions of the original experimental system sample. Here, from the liver of mice two celltypes are isolated. The liver isolation has the same protocol while cell type isolation has differing protocols. The respective files are expected to be on the same level as the Metadatasheet in the filesystem.

The next two segments ‘Preparation’ (Fig. 3D) and ‘Measurement’ (Fig. 3E) capture the information for sample preparation approaches and measurement techniques, respectively. The ‘Preparation’ segment holds the information about the process of the experimental system to the specimen that gets measured. The respective protocol can be selected from a predefined set of terms, such as common workflows or entering a filename in the designated comment field, as shown here. When there are subsamples present (Fig. 4B), information at segments’ secondary level is necessary, such as the number of subsamples per sample, their instances, replicates, and preparation information must be provided in a tabular format. The ‘Measurement’ segment requests details depending on the respective choice of the measurement technique (Fig. 3E). Note, that ‘used facility’ was an additional dependent key added upon the process of filling the Metadatasheet. The user can easily add further keys by entering the wanted key in both dependent fields sheet in respective column of Measurement type: ‘bulk_RNA_seq’ and specify its type of constraints, e.g., free-text, date or controlled vocabulary, within the ‘Validation’ sheet.

The final segment ‘DataFiles-Linkage’ (Fig. 3F), connects the measurement results with metadata. On the first level, one specifies whether raw or processed data is available. Raw data denotes the original machine-generated output, untouched by any processing, here the raw data are the .fastq files. At secondary levels, users would provide more details about their file naming system. Three options are provided: ‘ID contained in filename’, ‘single file for all’, and ‘CHANGES’. The options ‘ID contained in filename’ and ‘single file for all’ require the data to be positioned at the same level as the metadata document within a file system, whereby relative paths can be given. The option of ‘CHANGES’ (switching key:value pair to tabular form) allows user to define their unique naming system in the Measurement-Matching section. For processed data the procedure is required, and to be provided like the preparation protocol.

#### Example Measurement-Matching Section

The last but the most important step for Metadatasheet is the ‘measurement-matching’ section, which links the recorded metadata to the measurement data. This section involves an ID-specific metadata table to facilitate matching (Fig. 5). Here, the measurement for each replicate within a group requires a unique measurement ID. Given this ID and the group name (defined at top of Metadatasheet), one must be able to identify respective measurement. If there are subgroups or further subdivisions of samples, a table per division is expected. By design, the actual measurement happens at the last division stage, hence the measurement ID belongs to the last stage, as well. If available further personal IDs can be given on sample level, too.

**Figure 5.**
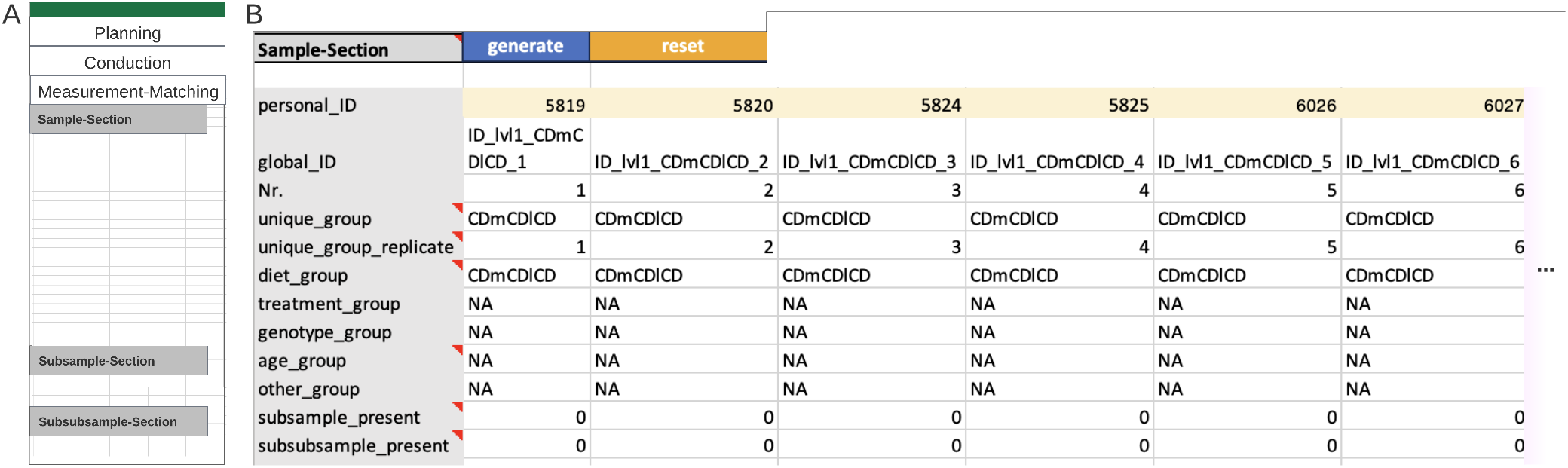
Example of an instance of the Measurement-Matching section. **(A)** Overview Measurement-Matching section. **(B)** An ID-specific metadata table example with the minimal number of required rows. The yellow marked cells hold measurement IDs (‘personal_ID’) required for the matching of metadata column with the respective measured data. ‘NA’ indicates non available information (‘Diet’ is the only comparison group specified). The last two rows indicate that neither subsamples nor subsubsamples are needed in this instance. The table is column cropped; based on previous final groups and given replicates a total of 30 columns are expected in the full table. Note that the full Metadatasheet of this example can be found in Supplementary Material.

The automatically generated ID-specific metadata table summarizes the preceding input of the user to easen the measurement to metadata matching. Hence, besides the default rows, the ID-specific metadata table will expand depending on inputs from the Conduction section. Expansion includes previously mentioned covariates and constants, along with any keys where the ‘CHANGES’ value was applied. The Measurement-Matching section overall ensures the flexibility tailored to capture information individually for each measured sample or division of such. Moreover, the arrangement of subsamples and subsubsamples clearly reveals any nested design, which is important for choosing appropriate statistics.

### Application of complete Metadatasheets

The availability of standardized Metadatasheets offers advantages to both individual users and the scientific community, ranging from the respective group to large-scale consortia.

The individual’s benefits from utilizing the Metadatasheet as a live document or central hub guides their data management for conducted or planned experiments. This approach simplifies the process of handing or taking over projects, as documentation follows a streamlined format, as opposed to each person maintaining individual data management methods. Furthermore, standardization plays a pivotal role in enabling the development of programs for analysis and processing, thanks to uniform input formats. A notable example is the provided conversion program that parses the Metadatasheet involving bulk-omics measurements to an R object. This SummarizedExperiment object^21^ itself is the standarized input for many Bioconductor based analysis^23^,^24^.

A group or consortia introducing the Metadatasheet will have access to multiple Metadatasheets. This in turn evokes the possibility for creation of a comprehensive database. Within this database, numerous sheets can be easily searched for specific information. To support this application, we have developed a dedicated, publicly accessible ontology for seamless integration of data into a custom database. Essentially, this database functions as a centralized knowledge hub, enabling swift access to available data, available specimen and planned experiments across groups. A database facilitates meta-analyses and aids in identifying gaps in the current local research landscape potentially discovering collaboration opportunities.

In summary, the application example showcases the Metadatasheet in practice. The use of Metadatasheets benefit individual users and the scientific community by streamlining data management and enabling program development.

## 3 Discussion

The developed metadata standard facilitates comprehensive recording of all relevant metadata for a broad spectrum of biomedical applications throughout the data-lifecycle. The standard’s implementation ensures efficient documentation of metadata and with a user-friendly design. The provided Metadata Workbook enriched with custom, open-source functionalities can be extended on various levels to adjust to additional setups.

The presented framework, encompasses two parts. The first part involved the iterative collection and organisation of keys, while the second part focused on the implementation of the user experience within the Metadata Workbook. During the collection phase, it became apparent that the specific set of keys varies enormously depending on the research groups. To address the high variability, we made adaptability of the Metadatasheet a priority. While the set of comparisons (‘comparison groups’) is tailored to our context, e.g. diet or temperature, the implementation is designed to be extensible ad-hoc. This means the Metadatasheet can be customized by specifying requested keys and adding experimental groups and measurement types, as well as expanding the controlled vocabulary. Moreover, a versatile comparison group labeled as ‘Others’ has been introduced. This ‘Others’ group adapts to any comparison scenario, not covered. Adding another ‘comparison group’ to the structure is also possible when adhering to the segments structural characteristics, only requiring additions to the provided Metadatasheet ontology.

The Metadatasheet has been implemented within a macro-enabled Microsoft Excel workbook. Despite the fact that Excel is not open-source, nor free it has several severe advantages. Its widespread availability, familiarity and standard-use within the biomedical research community makes it a valuable choice, especially when compared to custom standalone applications. Furthermore, most users are experienced Excel user, allowing for seamless integration of our proposed sheet into existing workflows. This immediate integration would not be as straightforward with open-source spreadsheet software like LibreOffice, also lacking required automation aspects. An online, browser-based, operating system independent approach, besides being accessible for everyone, violates the needs of sensitive data, particularly in cases involving unpublished studies. Recently, Microsoft has introduced Excel365, a browser-based software. However, our provided Metadata Workbook, requires adjustments to function within the Excel365 framework, as the used automation languages differ.

Metadata labels provide meaning to data, especially if keys and values are not only comprehensive but also interconnected, enabling cross-study comparisons. Providing metadata labels is commonly referred to as semantic interoperability, and it is considered a pivotal aspect of data management^25^. In order to attain semantic interoperability, there are domain-specific ontologies that establish meaningful connections between the labels of metadata. However, it is important to note that there is no single ontology that can comprehensively address the diverse requirements, even within a relatively homogeneous domain of investigation within a single consortium in the field of biomedical sciences. In fact, the choice of the appropriate ontology is far from straightforward and can vary for the same keys depending on the context. Pending ontology decisions might delay the recording of metadata, which in turn can lead to data loss. Involvement of inexperienced users, due to common high fluctuations of early-stage researchers, can further exacerbate the delay. Therefore, we have made the conscious choice, following our adaptability priority, to employ an extendable controlled vocabulary. This decision empowers biomedical researchers to directly and effortlessly record metadata without the need to immediately handle ontologies and their unavoidable complexities. While this decision will require additional retrospective annotation efforts to adhere to appropriate ontologies, it is manageable in contrast to retrospectively recovering metadata information that was never recorded. Our strategy prioritizes ease of initial data recording and acknowledges the practical challenges associated with ontology selection and application.

Ontologies enrich any set of collected metadata, therefore, we do not aim to discourage the use of ontologies. Integration of ontologies into the workflow could be facilitated by Metadata Annotation Services, such as RightField^8^ or OntoBee^26^. RightField is a standalone tool populating cells within a spreadsheet with ontology-based controlled vocabulary. OntoBee is a linked data server and can be used to query suitable ontologies and IDs given a keyword. Groups can enforce the partial or complete usage of ontology for keys in the Metadatasheet by leveraging on the option of group-specific validation and creating a tailored validation sheet.

We anticipate our proposed Metadatasheet accompanied with its implementation, the Metadata Workbook, being used for more than just data recording. Even in a partially filled state and at the start of a research cycle, the findability, accessibility, and interoperability provided by standardized Metadatasheets can speed up experiment preparation between groups, encourage effective specimen usage, and foster collaborations. The further planned deployment of the Metadatasheet and workbook includes adding export options, a database for Standard Operation Protocols, analyzing sets of collected metadata, and providing project monitoring tools. In conclusion, the framework leverages the widespread use of Excel, enabling comprehensive metadata documentation and improving the efficiency of data deposit on repositories. Our practical solution offers a user-friendly and sequential approach to manage metadata, thereby addressing the need for FAIR data in the field of biomedical science at intermediate stages during the data life cycle up to publication. We expect this to be of high relevance for a broad spectrum of biomedical researchers, and think that it can also be easily adapted to adjacent fields.

## Methods

### Metadata Workbook Structure

The proposed Metadatasheet is implemented within Microsoft Excel macro-enabled workbook, which consists out of multiple sheets with macros modules. The input sheet resembles the Metadatasheet. The other sheets hold the validation resources, the dependent fields for the differing experimental systems and measurement types, a plain Metadatasheet for reset, the repositories metadata standards, and additional resources for user guidance, such as a glossary. Input, validation, dependent fields and user guidance sheets are visible to the user, whereby only the input sheet is extensively editable by the user. Within validation and dependent fields sheets only blank cells can be filled.

The structure of the individual sheets ensures their functionality. An example is the validation sheet which holds per column the controlled vocabulary for a respective key. Each column starts with the three rows where the type of validation - freetext, date, DropDown or DropDown_M (multiple selection possible) - any specification in form of help text and the respective key is specified. The depended fields sheet is constructed in a similar manner. Here, the first two rows for each column determine the general category - measurement type or experimental system - as well as the specification from the controlled vocabulary set e.g. of mouse. After those specifications, the dependent keys are enumerated.

The input sheet and attached functionalities utilize different font faces as well as color cues for structuring, and segment specific automatised processes. All grey cells with bold font content signal different segments of each section. This provides a fine-grid structure. Italic font characterize boolean validation requests, hence expecting ‘yes’ or ‘no’. This does not only help for structure but also is done for performance reasons as just by checking font, actions can be precisely called.

### Custom add-on functionalities

The Workbook including VBA based macros was developed using Excel Version 16.77. The implementation is tested for use on both macOS (Ventura 13.5) and Windows (Windows 11) and respective variations of Microsoft Excel Version 16. The differences in Excel functionality between Windows and macOS influenced our implementation, such as bypassing ‘ActiveX-controls’ being not available on MacOS platforms.

The Metadata Workbook incorporates various functionalities organized into VBA modules. Users invoke actions by either actively pressing a button or upon input, which is a change of a cell within the input sheet. The latter allows for reactive updates. Reactivity functionality is directly attached to the input sheet unlike VBA modules. The Metadata Workbook key functionalities include a validation function, an insertion-of-dependent-keys function, and a reset-import function, which are further discussed in the following. Furthermore, the reactivity procedure evoked upon cell change is outlined.

The custom validation function leverages Excels Data-Validation feature. The feature checks predefined conditions for a given cell upon the user’s input, e.g. if the input values lies within a range of allowed values. If those values are of discrete nature one can display all possible values as a DropDown to the user. Our custom validation function populates Excels Data-Validation feature automatically, passing the appropriate data constraints to determine a valid input. An exception exists for all keys that allow multiple selections, marked in the validation sheet as type DropDown_M. To allow the selection of multiple items reactive functionalities had to be included. Any user values that fail validation are marked. To simplify searching within the DropDown list, the allowed values are automatically sorted alphabetically.

In the case of extensive controlled vocabulary or wish to tight constraints, users have the option to subset the main validation sheet. The subset sheet must be named ‘Validation_[Group]’, whereby ‘[Group]’ is to be replaced by the respective value to the requested key group. The structure of the subset sheet is expected to be the same as within the validation sheet. To use this predefined subset one has to choose ‘yes’ for ‘group specific?’ on top of the sheet.

The insertion functionalities handle the automatic dependent key insertion, inserting necessary keys dependent on the user’s choice of the experimental system and measurement type. Here, the subroutines conduct a search for a match with the user’s input within the ‘dependentFields’ sheet, retrieving the corresponding column with associated keys for insertion in the Metadatasheet. Note that dependent key sets can be extended by adding keys to the list, whereby additional keys subsequently need to be added to the validation sheet to provide constraints.

The reset/import function allows users to reset the sheet to its initial state or to a chosen template state. Two options are available upon pressing the ‘Reset’ button and displayed to the user with a pop-up window. The first option resets to a blank input sheet. The function deletes the current input sheet, copies a ‘ResetSheet’ and renames it to ‘Input’. The ‘ResetSheet’ has the same VBA-code as the ‘Input’ Sheet attached. The second option resets to a user chosen template. A template may be a previous complete Metadatasheet or a partially filled Metadatasheet. The inputs from the template sheet are copied upon a duplication of the ‘ResetSheet’ to retain reactivity-functionality. The duplication with the template’s input is renamed to ‘Input’. The original ‘ResetSheet’ is always hidden to prevent accidental deletion.

### Metadatasheet ontology creation

Our custom ontology was modelled by following a top-down approach using established tools in the realm of semantic web (cf. Protégé^27^ and accompanying tools), giving rise to a consistent contextual data model, logical data model and physical data model eventually leading to an integration of individuals (metadata samples) into a semantic database.

### Conversion program creation

The conversion program uses a completed Metadatasheet as input and checks for suitability of conversion based on the measurement type. If the type is one of ‘bulk-metabolomics’,’bulk-transcriptomics’ or ‘bulk-lipidomics’, the conversion starts. The Measurement-Matching section will be saved within ‘colData’-slot. The actual data matrix is identified guided by the Data File Linkage information. Given the personal ID and the given file measurement data is identified. Note, the location of the input Metadatasheet is seen as root and given filenames are expected as relative paths. If ‘single file for all’ is selected the filename given in the comment section is directly searched for. If nothing is found measurement data is searched for by the given extension in processed data and returned to user asking for clarification. The program is written in R.

## Data availability

The ontology needed to create a database upon a set of Metadatasheets (version 1.8.0) is available under the following link on Github https://github.com/stephanmg/metadata_ontology.

## Code availability

The Metadata Workbook and related content is freely available on Zenodo (free access upon publication - now reachable by provided link) and GitHub (https://github.com/LeaSeep/MetaDataFormat). The repository contains the macro-embedded Metadata Workbook, the isolated VBA scripts, as well as the converter to turn a Metadatasheet to a SummarizedExperiment Object.

## Acknowledgements

This work was supported by the Deutsche Forschungsgemeinschaft (DFG, German Research Foundation) under Germany’s Excellence Strategy (project IDs 390685813 - EXC 2047 and 390873048 - EXC 2151) and through Metaflammation, project ID 432325352 – SFB 1454 (L.Se., I.S., H.H., D.Ri., J.Y., T.B., K.S., R.K., S.K., E.M., D.W., E.L., F.M., A.Sch., J.H), BATenergy, project ID 450149205 - TRR 333 (S.G., A.S.A., S.H., M.M., D.Ra., J.Be., D.W., A.T., V.P., K.K., A.P., H.W., L.Sch., T.F., W. K. F., M.K., J.H), the Research Unit “Deciphering the role of primary ciliary dynamics in tissue organisation and function”, Project-ID 503306912 - FOR5547 (D.W., E.M.), and SEPAN, project ID 458597554 (L.Se.), and by the University of Bonn via the Schlegel professorship to J.H.. E.M. is supported by the European Research Council (ERC) under the European Union’s Horizon 2020 research and innovation program (Grant Agreement No. 851257). W.K.F. is further supported by the DFG (FE 1159/6-1, FE 1159/5-1, DFG FE 1159/2-1), by the European Research Council (ERC, under the European Union’s Horizon Europe research and innovation program; Grant Agreement No. 101080302) and by grants from the Gabriele Hedwig Danielewski foundation and the Else Kroener Fresenius Foundation. A.T. is supported by the Gabriele Hedwig Danielewski foundation. A.K.G. is supported by Medical Faculty, University of Bonn, BONFOR grants 2018-1A-05, 2019-2-07, 2020-5-01.

We thank all members, including associated, of the SFB Metaflammation and TRR BATenergy for the iterative discussions and their input throughout.

## Author contributions statement

J.H. and S.G. conceived the concept. L.Se. implemented and extended the Metadatasheet and created the Metadata Workbook. T.B., M.K., J.Br., A.St. tested and provided feedback on initial version of the Metadatasheet. I.S., D.Ra., M.M., S.H., M.L., K.H., D. Ri, K.S., R.K., H.H., J.Y., S.K., J. Be., A.T., V.P., A.S.A., D.T., K.K., Y.L., M.B., A.K.G., T.F., H.W., M.K., W.K.F., L. Sch., F.M., A. Sch., E.M., D.W. provided in-depth feedback to the Metadatasheet and the Metadata Workbook and contributed to the showcases. E.L. and A.P. lead the discussion rounds as representatives of the consortia. L.Se. and J.H. wrote the first draft of the manuscript. All authors reviewed the manuscript.

## Competing interests

The authors declare no competing interests.

## Figures & Tables

Figures placed at suitable positions within the manuscript for revision.

## Notes

https://zenodo.org/records/10278069?token=eyJhbGciOiJIUzUxMiJ9.eyJpZCI6IjhlNjYxNTgwLTcxOTAtNDI2OC1hNDY4LWIxYjYxMzE2OTg1NSIsImRhdGEiOnt9LCJyYW5kb20iOiIyODdiYzgyMDE3YWMxYWQ3MWQ3NmY3YTJkNTg1NjRkNiJ9.xYfLnm1sckepdmbNmsVhhwsAWwR0vJEKRcwoTiT1oks4kdPcI29Q-ikQkg0xz-XoEX2y7YORvJFXuB_AN9LszA

https://github.com/LeaSeep/MetaDataFormat

https://github.com/stephanmg/metadata_ontology

### Competing Interest Statement

The authors have declared no competing interest.

