## Supplementary Material for "From Planning Stage To FAIR Data: A Practical Metadatasheet For Biomedical Scientists"

### 4 Supplementary Material

#### Set of example Metadatasheets

In this section, several complete showcases are briefly described. Respective specific features highlighting a certain aspect of the proposed Metadatasheet are further discussed. You can find all complete Metadatasheets on Zenodo as well as all accompanied protocols files, if publicly available. As the Metadatasheet is to be filled alongside the data-lifecycle, majority of the measurement data as well as some protocols are under non-disclosure. Note, that the provided folder structure within each showcase is also as expected by the Metadatasheet:

**Bulk RNAseq from mouse samples with a one-fold experimental design:**

This example is associated with a publicly available dataset generated by Elvira Mass research group, accessible through the GEO Series accession number GSE237408. This dataset was created as part of a project aimed at understanding how maternal obesity affects the development of Kupffer cells and its implications for fatty liver disease within their offspring. The detailed example is part of a collection and linked via the value given in the 'title' key.

The data shown here involves mouse as the experimental system and bulk-RNA sequencing as the measurement technique. We compared different diets, which is a part of a more complex design involving maternal diet, maternal diet during lactation, and offspring diet. Respective new terms for diet controlled vocabulary were introduced. While we could have chosen a two-fold design utilizing comparison group 'others', combining maternal diet and offspring diet, we opted for keeping all diet related groups within the same comparison group to enhance the findability of the data.

Note that 'Kupffer Cells' were a constant factor in this study. There's a similar experimental design for Hepatocytes in the collection. Our Metadatasheet could handle both within the same Metadatasheet by using sub-samples, as indicated in Figure 4 in the main manuscript. However, since this Metadatasheet was also submitting to GEO independently, we decided to keep the same setup as in the repository for consistency.

Lastly, the specified procedure used in this study is taken exemplary from the controlled vocabulary. This specification is accompanied by the relevant parts of the manuscript, which are stored on a consortium-wide file location (protocol hub) under selected value for the procedure. For completion the protocol is added as if it would have been specified as filename. In this example, we don't have any covariates to consider, so the Measurement-Matching section contains only default rows.

**Bulk metabolomics from cell-line samples with a one-fold experimental design:**

This example is associated with the public available dataset generated within Karsten Hiller group. The dataset was created as part of a project aimed to analyse the glucose metabolism within human monocyte derived macrophages in response to LPS stimulation.

The data shown here involves here human-derived as experimental system. The measurement technique is 13C labelling, done by the Department of Bioinformatics and Biochemistry, which as no special appointed measreument-type but is associated with the type ’bulk-metabolomics’ and associated dependent keys. The comparison group involves LPS treatment, at differing timepoints, which are idependent form each other, hence no nesting was used.

Further, the specified protocols are provided as additional file, which filename is specified in the Metadatasheet.

**Bulk proteomics from mouse samples with a one-fold experimental design:**

This example is associated with the data obtained from Anne-Katrin Gellner. The dataset was created as part of the project to decipher the role of stress vulnerability in the disruption of motor cortical neuroplasticity.[28].

The data shown here involves mice as experimental system which are exposed to a certain type of stress or not. The measurement technique is proteomics analysing the mice’s cerebrospinal fluid (CSF). After stress-treatment mice were classified as different phenotypes based on behavioral tests. Note, that these phenotypes are not the comparison groups as this assignment is made on additional data and not set *a priori*. Moreover, not all samples taken from mice have associated data due to different ’reason of drop out’, hence provided processed data encompasses a subset of provided mice IDs.

**Bulk proteomics from human-derived samples with a one-fold experimental design:**

This example is associated with unpublished data and results generated with Alexander Pfeifers group. The dataset encompassed human-derived material as experimental system, whereby the studied subjects differ in their sporting activities. Upon consultation the on ’human-derived’ keys did not encompass yet the key ’cellular component’, which was needed as extra-cellular vesicles where studied. Leveraging on the Metadata Workbook capabilities, we added immediately the missing dependent key to the lists within the ’dependent fields’ sheet at the column of human-derived as experimental system. Further, we added a set of controlled vocabulary, by adding the new introduced key ’cellular component’ to the ’validation’ sheet, and providing a list of plausible terms. After this we could further fill out the Metadatasheet, having only minor delay in recording.

Further, data was recorded ’pre’ and ’post’ from the same individual yielding a nested design. Hence. the time-point of measurement defines the subsamples.

Note, within the shown example, certain variables where left out due to confidentiality as this is an unpublished study. The provided measurement data outlines the structure of the data.

**16S rRNA from rat samples with a one-fold experimental design:**

This example is associated with a public available 16S rRNA dataset generated within the group of Wiebke K. Fenske. The dataset was created as part of a project aimed at understanding the relationship of bariatric surgery as the most effective therapy for weight loss and type 2 diabetes remission and the role of gut microbiota in mediating the surgery’s metabolic benefits[29].

The experimental system studied is ’rat’ which as been not been part of the experimental system collection upon consultation. As in previous showcases described, we added the missing experimental system to the list, providing requested keys and their respective controlled vocabulary, where necessary. The performed intervention was for now added to the ambiguous ’other’ group, allowing free text to describe each instance. As the ’other’ group might become really diverse, a group or a consortium might want to bundle a homogeneous group and create a new comparison group, analogous to the already present comparison groups, e.g., ’diet’ or ’temperature’. The introduction of a new comparison group, upon ontology adjustment, will help others to more easily find related experimental set ups among a set of Metadatasheets.

The analysis of gut microbiota composition by 16S rRNA sequencing was conducted by the company BGI (Hong Kong, China). The provided ’personal_IDs’ within the Measurement-Matching section matches with the public available dataset associated with the BioProject: PRJNA735921.

**Indirect calorimetry from mouse samples with a one-fold but nested experimental design:**

This example is associated with a public available dataset generated within the group of Alexander Pfeifer. The dataset was created to study the energy expenditure via extracellular inosine within apoptotic brown adipose tissue[30].

The studied experimental system was ’mouse’ as living animals. Hence, several further keys within the ’experimental system’ segment, such as tissue type, are not applicable and marked as such. The contrasting setting was the genoytpe of the respective mice. For the actual measurement (indirect calorimetry) the mice were separated into single cages. This is noted within the Time-Dependence-Segment as ’interuption-type continued’, as the genotype has not changed but the before actual measurement animals were relocated, which can result in stress. Indirect calorimetry measurements are obtained over the timecourse of day, of a single mouse. Hence, timepoints are subsamples and present a nested design. Weight was recorded as covariate and only measured once for each mouse and not each timepoint. By design, covariates are captured at lowest level, leading to duplicated weights measure.

Processed data is provided as Excel table, including statistical testing. Note, that the ID matching involves Sample IDs as well as Subsample IDs. This specialty is defined within the comment key of the Data File Linkage segment.

**Single-cell RNA-seq from mouse with a two-fold and nested experimental design:**

This example is associated with unpublished data and results generated within the group of Marc Beyer. The dataset was created to study the transcriptional programming of adipose tissue macrophages during metaflammation. As comparison groups there diet, including 4 different diet regimes, and genotype, including 2 different genotypes. The Timeline-Dependence segment was used to further specify the different diet regimes, as they differed in time they received high fat diet, Measurements where taken on three tissues, specified using ’subsamples’. As measurement type ’scRNA-Seq’ was selected, however, data is under non-disclosure. Example IDs are provided in respective fields.

**Single-nucleus RNA-seq from mouse with a one-fold but nested experimental design:**

This example is associated with unpublished data and results generated within the group of Tobias Fromme. The dataset was created to study the enhanced beige adipocyte detection by single nulcei RNAseq within a luciferase reporter mice. The comparison group is ’genotype’. Measurements where taken on two differing tissues, specified using ’subsamples’. As measurement type 'scRNA-Seq' was selected, as single-cell and single-nucleus measurements differ in the measured component but not with respect to the machine or other herein measurement-dependent keys. To clarify the difference, the preparation comment field and/or the measurement-dependent comment field should be used.

Provided data encompasses the raw FASTQ files for each subsample, as well as the processed data from cellranger software, which is commonly used as input for bioinformatic analysis. Note, to reduce data storage, a picture of actual raw data filenames (despite actual files) is provided. Also, a screenshot of processed data folders is provided. Those folders are appropriate named and hold multiple processed files.

**Bulk lipidomics from mouse samples with a two-fold and nested experimental design:**

This examples is associated with unpublished data, protocols and results generated within the group of Ludger Scheja. Here, two comparison groups are present, 'diet' and 'genotype' rendering the experimental design as two-fold. Additionally, two tissue types (two subsamples) where extracted and measured from each sample.

In this study the weight of the mice where recorded as covariate, as specified in the covariate/consants section. Within the Metadata Workbook this covariate is automatically added to the subsamples table within the Measurement-Matching section. As the weight of the mouse and not the weight of the subsamples was recorded, the weight matches the sample match and hence is duplicated on subsample level.

Note, that data as well as protocol are not yet available to the public.

**Lipolysis measurements from cell-line samples with a one-fold but nested experimental design:**

This example is associated with unpublished data and generated within the group of Alexander Pfeifer. The dataset was created aimed at unterstanding GPCR induced lipolysis in adipocytes.

The experimental system 'cell-line' was studied, measuring the lipolysis within the supernatant of the cells. The taken cells where contrasted with respect to the comparison group 'treatment', having nine instances. Each of those instances where measured with three biological and 2 technical replicates. The lipolysis measurements are taken within a well plate.

The lipolysis measurement type was not part of the measurement types upon consultation, therefore added as described in previous showcases. The raw measurements are provided as single Excel sheet, whereby the IDs refer to the respective well-location and the unique experiment identifier. The processed data is provided as Excel Workbook, including necessary calculations to get to the final lipolysis read out.

**UPLC-UV measurements from cell-line samples with a one-fold but nested experimental design:**

This instance is affiliated with data generated within the group of Alexander Pfeifer. The investigation focused on unraveling the purines release mechanism from brown adipose tissue, within the experimental system mice. The study encompassed two distinct treatments and four 'other' groups, here involving different types of virus infections. The experimental timeline of virus treatment was detailed within the Time-Dependence segment. UPLC-UV measurements, which were not originally included in the initial set of measurements, were incorporated into the Metadata Workbook leveraging its adaptability.

Within this study, two technical replicates were acquired for each biological sample, and all samples were originally prepared on a well plate. The provided dataset includes both the raw data and its processing, including a measured standard curve for each purine. It is important to note that standard measurements are not documented in the Metadatasheet records. To exemplify the integration of well-plate design into the Metadatasheet, the provided data includes the original plate design.

**FRET measurements from cell-line samples with a one-fold, timeseries experimental design:**

This example is associated with unpublished data generated within the group of Alexander Pfeifer. The dataset was created to investigate real-time cAMP and PDE dynamics in murine brown adipocytes.

The experimental system 'cell-line' was studied, taking Fluorescence Resonance Energy Transfer (FRET) images. The studied cells were consecutive treated with different drugs. Hence the comparison group 'treatment' was chosen and a instance of 'timeseries' was specified. This 'timeseries' instances was further annotated with time-dependent information within the 'Time-Dependence-Timeline'-segment.

The provided raw measurements are the FRET images which are stored within a tiff-series, with consecutive frames (not provided within Supplementary). One could have noted each frame as subsample (nested design). However, this is not how the raw data is actually provided, potentially introducing errors through dissection. Note, here only one example time-series is included, associated, processed intensity read-out is provided.

**Histology from mouse samples with a one-fold experimental design:**

This examples is associates with unpublished data generated within the group of Dagmar Wachten. The dataset was created to histologically analysis white adipose tissue from a genetically modified mice.

Here, both comparison groups received high fat diet while their genotype differed. For the two groups the weight was captured as covariate, whereby the high fat diet-fed duration was held constant as well as only male mice were studied.

Mentioned protocols are handed along the respective Metadatasheet. An example filename for a real image is also provided.
